# Parkinsonian striatal acetylcholine dynamics are refractory to L-DOPA treatment

**DOI:** 10.1101/2025.11.06.686401

**Authors:** Eva V. Potjer, Xunhui Wu, Allison N. Kane, Jones G. Parker

## Abstract

**Background:** Fluctuations in striatal dopamine (DA) are interdependent with fluctuations in acetylcholine (ACh), both of which are thought to be important for motor function and dysfunction in Parkinson’s disease (PD).

**Objectives:** Determine how ACh dynamics are altered by dopamine under normal and parkinsonian conditions following acute and chronic L-DOPA treatment.

**Methods:** We used fiber photometry to record fluorescent DA and ACh sensors in the dorsomedial striatum (DMS) of normal mice treated with vehicle or amphetamine and in the dorsolateral striatum (DLS) of mice with unilateral substantia nigra (SNc) DA neuron lesions (using 6-OHDA) subjected to an L-DOPA-induced dyskinesia (LID) protocol.

**Results:** Under normal conditions, ACh exhibited transient increases followed by pauses and fluctuated at ∼1-Hz frequency in an anti-correlated manner to DA. Amphetamine treatment reduced the frequency of ACh fluctuations and amplitude of ACh transients while prolonging after-transient pauses. In 6-OHDA-lesioned mice, ACh dynamics oscillated at a higher frequency, and ACh events had reduced peaks and pauses. L-DOPA increased DA and, after repeated treatments, also reduced ACh. However, L-DOPA neither restored the anti-correlation between DA and ACh nor normalized the reduced ACh transient and pause amplitudes or the increased frequency of ACh fluctuations. In extended recordings during the transition to the L-DOPA OFF state, ACh remained low as DA diminished, and ACh transient and pause amplitudes remained suppressed, and the oscillatory frequency elevated.

**Conclusions:** Dopamine depletion transforms striatal ACh dynamics and L-DOPA fails to normalize these dynamics. Because these dynamics are unresponsive to treatment, new therapies targeting them may offer symptomatic improvements for PD and LID.

**Graphical Abstract:** 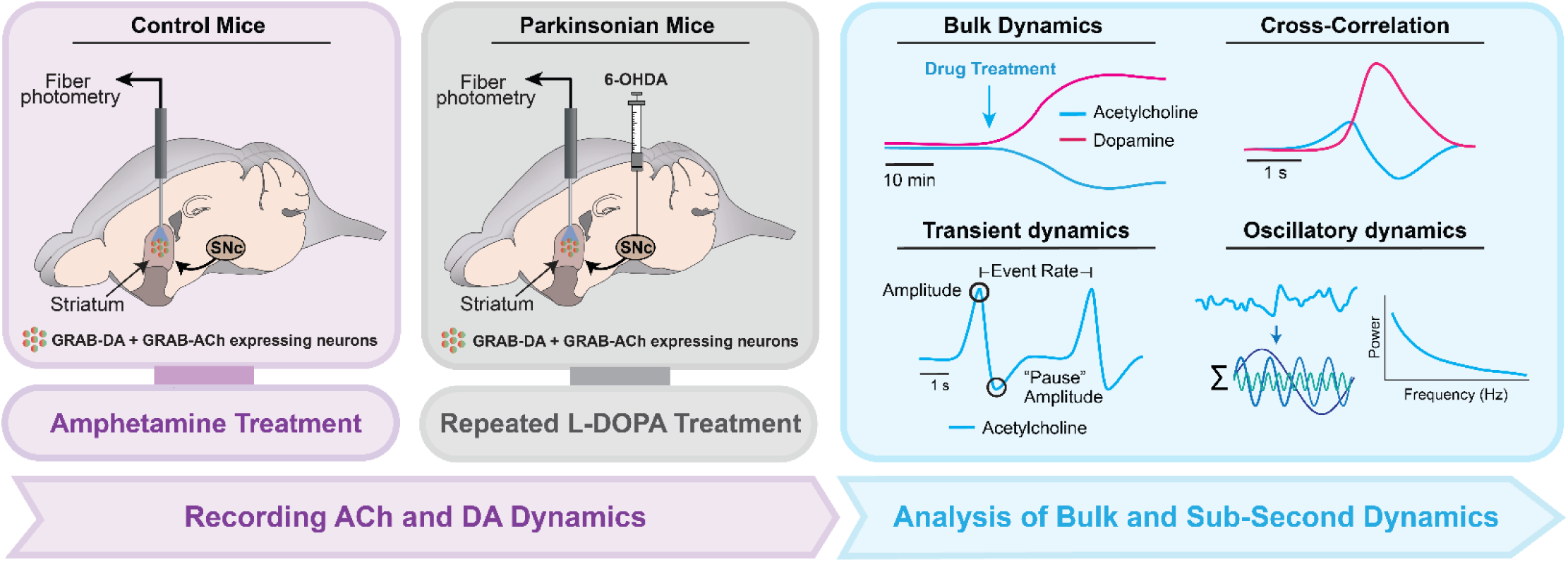

We used dual-wavelength fiber photometry to simultaneously record dopamine and acetylcholine dynamics in the dorsal striatum. We recorded these dynamics in control mice before and after treatment with the dopamine releaser amphetamine and in hemiparkinsonian mice before after treatment with the dopamine pre-cursor L-DOPA. We asked how these experimental manipulations alter the bulk signaling of dopamine and acetylcholine, their temporal relationship, and the sub-second dynamics of acetylcholine signaling.

## Introduction

The modulation of neural activity in the striatum, the principal input stage of the basal ganglia, is crucial for motor function. Within the striatum, dopamine (DA) and acetylcholine (ACh) levels are both among the highest of any brain region^1,2^. In Parkinson’s disease (PD), the loss of substantia nigra (SNc) dopamine neurons diminishes DA in the striatum, but also alters striatal ACh^3^. This abrogation of the normal balance between striatal DA and ACh is predicted to impair movement by dysregulating the activity of D1 and D2 DA receptor-expressing striatal output neurons (SPNs). For instance, D1-SPNs express inhibitory M4 ACh receptors, through which increased ACh may counter-modulate excitatory D1 receptor signaling. Likewise, D2-SPNs express excitatory M1 ACh receptors, through which increased ACh may counter- modulate inhibitory D2 receptor signaling^4,5^. In addition to the overall balance in the levels of DA and ACh in the striatum, the relative timing of fluctuations in DA and ACh are thought to subserve important synaptic plasticity rules in D1-SPNs and D2-SPNs^6,7^.

While DA replacement with L-DOPA is highly effective for PD, how it influences ACh dynamics is not fully understood. Canonically, DA and ACh dynamics are thought to have an inverse relationship in the striatum. Consistent with this idea, L-DOPA treatment reduces overall ACh levels in the depleted hemisphere of hemiparkinsonian mice^8^. However, recent advances using *in vivo* imaging with optical sensors are revealing a more intricate relationship between striatal DA and ACh^9–15^. For example, striatal ACh interneurons and DA axons co-activate during movement onset^11,16^. Moreover, the magnitude of the decrease of “pauses” following transient ACh increases is scaled by DA^12,13^. DA and ACh levels also spontaneously fluctuate at a frequency of ∼2 Hz in an anti-correlated manner, and both exhibit wave-like spatiotemporal dynamics^14,15,17^.

Given their complex interrelationship, the loss of DA in PD could transform striatal ACh in a myriad of ways to exacerbate motor symptoms. It is notable that some of the earliest treatments for symptomatic treatments for PD were anticholinergic agents^18^. In particular, blocking M1 or M4 acetylcholine receptors has antiparkinsonian effects, potentially through actions in the striatum^19^. While anti- cholinergic drugs are not as effective as L-DOPA, the emergence of L-DOPA-induced dyskinesia (LID) after years of treatment continues to be a major therapeutic limitation. Aberrant ACh dynamics may also contribute to LID, as enhancing M4 receptor signaling mitigates chronic L-DOPA treatment-induced abnormal involuntary movements and aberrant synaptic plasticity in the striatum of parkinsonian mice^4,19^. Altogether, these findings suggest that better understanding how ACh dynamics become altered in the context of PD could yield new insights and novel therapeutic strategies.

To determine these dynamics, we combined a red GRAB-DA sensor and a green GRAB-ACh sensor with fiber photometry to simultaneously record DA and ACh dynamics in the dorsal striatum. We determined their dynamics in control mice under normal and hyperdopaminergic conditions and in parkinsonian mice with and without L-DOPA treatment. We found that driving dopamine release via amphetamine treatment reduced the peaks of ACh transients, shallowed, but prolonged post-transient ACh pauses, and slightly reduced the frequency of spontaneous ACh fluctuations and their cross-correlation to DA.

Dopamine depletion also reduced ACh transient peak amplitudes and markedly diminished the pause in ACh after each transient but increased the oscillatory frequency of ACh and abolished its inverse cross-correlation to dopamine. L-DOPA treatment increased bulk dopamine but exacerbated the effects of dopamine depletion on ACh transient dynamics and did not normalize its oscillatory dynamics. After the induction of LID, L-DOPA treatment caused a greater suppression of bulk ACh and a reduced ACh transient frequency. During transition to the L-DOPA OFF state, the L-DOPA-induced decreases in bulk ACh and ACh transient amplitude persisted, while the reduced frequency of ACh events and altered Ach oscillatory dynamics returned to pre-treatment levels. Overall, we found that dopamine depletion alters ACh dynamics and that L-DOPA treatment failed to normalize these dynamics, exacerbating some of them. Our findings provide newfound alterations in ACh signaling that may be therapeutically targeted to ameliorate the symptoms of PD and LID.

## Methods

### Animals

All mice were housed and handled according to guidelines approved by the Northwestern University Animal Care and Use Committee. We used C57BL6/J mice (Jax #000664) that were 8–12 weeks at the start of experimental testing.

### Virus injections

We anesthetized mice with isoflurane (2% in O2) and stereotaxically injected virus at a rate of 250 nL·min^-1^), using a calibrated glass micropipette (Drummond Scientific Company, Broomall, PA) pulled on a P- 97 Sutter Instruments (Novato, CA). For the mice used for un-lesioned recordings, we injected into the DMS (Anteroposterior (AP): 0.4, Mediolateral (ML): 0.6, and Dorsoventral (DV): -3.3 mm from bregma). For the mice that underwent 6-OHDA lesions, we injected into the DLS (AP: 0.25, ML: 2.5, and DV: - 3.4 mm from bregma). We adjusted the AP coordinates to account for individual differences in the bregma-lambda distance according to the following: adjusted AP = AP ÷ 4.21 × bregma-lambda distance (in mm). We injected mice in both cohorts with a 700 nL mixture of AAV2/9-hSyn-AAV-hSyn-rDA3m- WPRE-PA (1.00 × 10^-12^ vg·mL^-1^; BioHippo) and AAV2/9-hSyn-AAV-hSyn-gAch4m (3.37 × 10^11^ vg·mL^-1^; BioHippo). After the injection, we left the syringe in place for 10 min, then slowly withdrew the syringe. For the mice used for un-lesioned recordings, we performed the optic fiber implant at the site of virus injection in the same surgery, securing the fiber as described below. For mice that underwent 6- OHDA lesions, we sutured the scalp, injected analgesic (Buprenorphine SR; 1 mg·kg^-1^) and allowed the mice to recover for at least one week prior to subsequent surgery.

### 6-OHDA Infusions

We anesthetized mice with isoflurane (2% in O2) and injected mice with desipramine (25 mg·kg^−1^; i.p.) 30 min before 6-OHDA infusion. We then stereotaxically injected 6-OHDA (4 μg·μl^−1^ in saline) at a rate of 100 nL·min^-1^ into the SNc using a microsyringe with a 33-gauge beveled tip needle (WPI; Nanofil). We sequentially injected 6-OHDA at three sites (AP: -3.5, ML: 1.25 from bregma, DV: -4.2, -4.0, and - 3.8 mm from dura), with adjustment to AP coordinates for individual bregma-lambda distance. After each injection, we left the syringe in place for 10 min, then slowly withdrew it to the next site or out of the brain after the final injection. We then sutured the scalp, injected analgesic (Buprenorphine; 1 mg·kg^-1^), and subcutaneously injected 1 ml of Ringers Solution to prevent dehydration. We closely monitored the mice for 3 days following surgery, providing moistened food and a heating pad.

### Fiber optic Implant

For our photometry experiments with 6-OHDA lesioned mice, we anesthetized virus-injected and lesioned mice with isoflurane (2% in O2) and used a 0.5-mm diameter drill bit to create a craniotomy (AP: 0.25 and ML: 2.5 mm from bregma) for implanting an optical fiber (MFC_400/430-0.48_4mm_MF1.25_FLT; Doric). We used a 0.5-mm diameter drill bit to drill four additional small holes at spatially distributed locations for insertion of four anchoring skull screws (000-120 x 5/64 SLOTTED FILLISTER; Antrin miniature specialties). We implanted the optical fiber at DV: -3.4 mm from bregma and we applied Metabond (C&B) to the skull then used dental acrylic (Coltene) to fix the fiber along with a custom stain- less-steel head-plate (Laser Alliance) for head-fixing mice during attachment and release of the optic fiber to the patch cord. We injected analgesic (Buprenorphine SR; 1 mg·kg^-1^) and allowed the mice to recover for around 1 week prior to recording.

### Drugs

We injected all drugs subcutaneously at a volume of 10 mL·kg^−1^. We dissolved D-amphetamine hemisulfate (2.5 mg·kg^−1^), donepezil (2 mg·kg^−1^), and L-DOPA (6 or 12 mg·kg^−1^) in saline (0.9% NaCl). To induce LID, we administered two doses of 6 mg kg^−1^ L-DOPA then three 12 mg·kg^−1^ L-DOPA every other day, two weeks after the 6-OHDA lesion. We administered additional 6 and 12 mg· kg^−1^ doses of L-DOPA on separate days for the final two photometry recordings. For all L-DOPA injections, we co-administered benserazide (12 mg·kg^−1^ i.p.) to inhibit peripheral L-DOPA degradation *in vivo*.

### Fiber Photometry Recordings

Prior to recordings, each mouse’s optic fiber implant was cleaned with 70% ethanol and tethered to a low-autofluorescence patch cord (MFP_400/430/1100_2m_LAF; Doric) by wrapping parafilm around the connection point to optic fiber and then connected to a commutator (Doric). We used a controller and LED sources (Thorlabs) with miniature filter cubes (FMC6_AE(400-410)_E1(460-490)_F1(500-540)_E2(550- 580)_F2(600-680)_S; Doric) to deliver ∼30–70 µW from the optic fiber tip of green (565 nm), blue (470 nm) and UV (405 nm) light modulated at 530, 330, and 210 Hz, respectively. The GRAB-ACh, GRAB-DA, and isosbestic fluorescent signals were collected through the same patch cords coupled to femtowatt photoreceivers (2151; Newport). We used a commercial system and software (RZ10x and Synapse; TDT) to sample at 1017 Hz, de-modulate, and lock-in amplify these signals. The signals were demodulated and amplified by the processor. Data was collected using the Synapse software and analyzed with custom MATLAB code. Prior to recordings, each mouse’s optic fiber implant was cleaned with 70% ethanol and the patch cord was secured by wrapping parafilm around the connection point to the optic fiber. For WT mice photometry recordings, we recorded for an initial 20 minutes to habituate the mice, then administered vehicle, recorded fluorescence for 20 min, then subcutaneously injected Amphetamine (2.5 mg·kg^−1^) or Donepezil (2 mg·kg^−1^), and recorded fluorescence for 40 min. For lesioned mice photometry recordings, the same procedure was followed except we injected L-DOPA (6 or 12 mg·kg^−1^), and subsequently recorded for 40 minutes for the first 4 recordings, and 180 minutes, toggling 5 minutes ON and 5 minutes OFF, for the final recording.

### Photometry Pre-Processing

Traces were converted from the TDT to MATLAB using the TDT2mat function provided by TDT. Next, the first and last 30 seconds of each recording were removed. To correct for photobleaching, the vehicle and drug period traces were concatenated and fit to the last 40% of the vehicle period trace using the fit MATLAB function with the exp1 model. For analysis of bulk dynamics, the trace was normalized by computing Δ*F/F* of the trace by subtracting and dividing the mean of the vehicle period from the full trace, multiplying by 100 and downsampling to 1-minute bins. To analyze transient dynamics, the traces were normalized by calculating the Δ*F/F* in 5 second windows along the full trace and down-sampled to 30 Hz.

### Transient Dynamics Analysis

#### Event Detection and Quantification

After processing and downsampling the traces, we identified the individual ‘events’ in the bulk-fluorescence trace using a threshold-crossing algorithm^20^. Within 60-second windows, we reduced any noise by subtracting a median-filtered version of the signal (using a 40-second sliding window) and then calculated the standard deviation (s.d.). We identified any peaks that were ≥ 1.5 s.d. while enforcing a minimum inter-event time of > 1.5 s. We determined the time of each GRAB-DA or GRAB-ACh event as the temporal midpoint of the time of each event’s fluorescence peak. For visualization, we averaged 1-second trace segments aligned to the detected event peaks (**Fig. 1C, D**). We used the same detected event indices to quantify peak amplitudes using the Δ*F/F* values calculated in 5-second windows. For each ACh event, we determined whether a DA event also occurred within 2 s after the ACh event peak. Events meeting this criterion were labeled as “with DA” and events without a subsequently occurring DA event as “with- out DA”.

**Figure 1.**
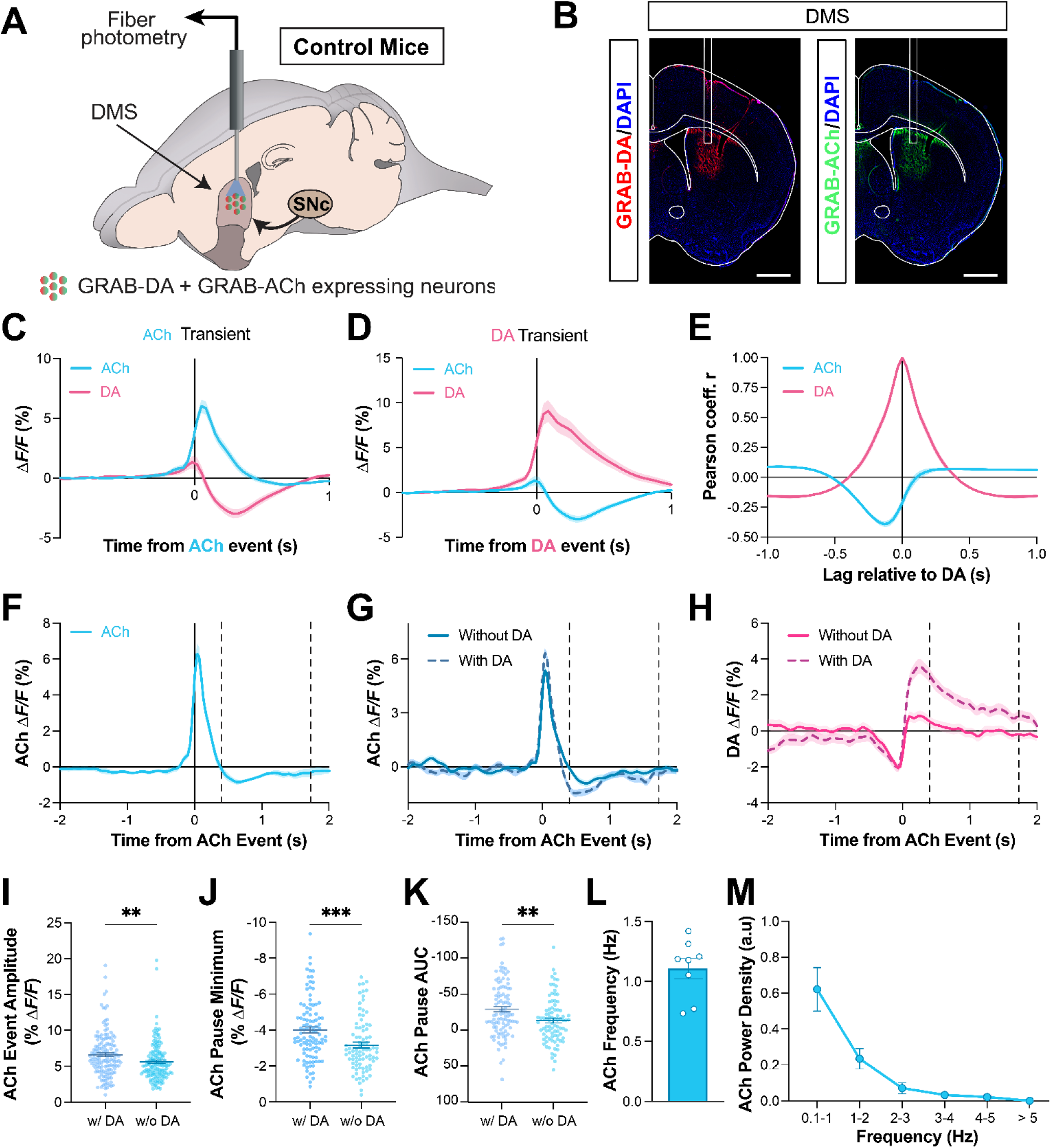
Measuring simultaneous striatal dopamine and acetylcholine dynamics. **(A)** We used an fiber optic to measure DA and ACh signaling via virally expressed GRAB-DA and GRAB-ACh in the DMS **(B)** GRAB-ACh and GRAB-DA expression in the DMS of a representative mouse (scale bar: 1 mm). **(C)** Mean ± s.e.m. GRAB-DA and GRAB-ACh fluorescence aligned to GRAB-ACh event peaks. **(D)** Mean ± s.e.m. GRAB-DA and GRAB-ACh signals aligned to GRAB-DA event peaks. **(E)** Mean ± s.e.m. cross-correlation between ACh (*blue*) and DA signals and DA signal autocorrelation (*pink*) **(F)** Mean ± s.e.m. GRAB-ACh event-triggered fluorescence. Vertical dashed lines denote the window over which we quantified GRAB-ACh pause metrics for each grouping in **J**, **K**. **(G)** Mean ± s.e.m. GRAB-ACh event-triggered fluorescence separated by events that occurred with or without a GRAB-DA event within the subsequent 2 s. **(H)** Mean ± s.e.m. GRAB-DA fluorescence corresponding to the separate groups in **G**. **(I–K)** Mean ± s.e.m GRAB-ACh fluorescence peak amplitudes, **I**, post-peak pause minima, **J**, and post-peak pause areas under the curve (AUC), **K**, with and without a subsequent GRAB-DA event (\*\**P* < 0.01 and \*\*\**P* < 0.001; unpaired t-test). **(L)** Mean ± s.e.m. weighted mean of GRAB-ACh fluorescence signal FFT frequency. **(M)** Mean ± s.e.m. power spectral density of the GRAB-ACh fluorescent signal in 1-Hz bins. Data in **C–F**, **L**, and **M** are the average of *N* = 8 control mice. Data in **G–K** represent *N* = 143 events of each category from of *N* = 8 control mice.

To quantify the pause in ACh signaling following a transient event, we defined a pause window to compute the pause minima and AUC values using the average x-axis crossing times after the peak of the average, event-triggered Δ*F/F* traces for GRAB-ACh. For comparisons between vehicle and amphetamine treatments, we adjusted the start and end of the pause window based on each condition’s average x-axis crossing times. For comparisons between “with DA” and “without DA”, and control vs. lesioned or L-DOPA–treated mice, we used the pause window determined from the vehicle treatment period of control mice (**Fig. 1F**).

#### Cross-Correlation

To assess the temporal correlation between GRAB-DA and GRAB-ACh signals, we computed the Pearson cross-correlation coefficient using MATLAB’s “xcorr” function. Cross-correlation was performed in 2- second windows across the full trace, with each window normalized to its own mean to control for baseline differences.

#### Frequency Analysis

To analyze the frequency components of the GRAB-DA and GRAB-ACh signals, we applied a Fourier transform to the raw, unfiltered, and non-downsampled signals from both vehicle and drug conditions. We computed the FFT using MATLAB’s “fft” function and extracted a single-sided power spectrum. Prior to transformation, we subtracted from each trace its mean to remove DC offset at 0 Hz.

To normalize all the power spectra and mitigate noise contributions, we subtracted the power at 100 Hz and divided by the power at 0.01 Hz. We then calculated a weighted mean frequency (weighted by power) to compare enriched frequency components across conditions. To normalize the power density spectra, we divided the spectra by the sum of power values between 0.1–10 Hz and averaged across 1-Hz frequency bins.

### Histology

After all photometry experiments, we euthanized and intracardially perfused mice with PBS followed by a 4% solution of paraformaldehyde in PBS. After perfusion, we extracted the brains and placed them in 4% paraformaldehyde for 1–3 days. We sliced 50-µM-thick coronal sections of the striatum and SNc from the fixed-brain tissue using a vibratome (Leica VT1000s). For immunostaining the SNc, we used α-Tyrosine Hydroxylase (1:500; Aves TYH) primary antibodies and Alexa 594 conjugated secondary antibodies (1:500; Jackson Immunoresearch #703-546-155 or #711-546-152 with #715-586-150, respectively) to verify unilateral lesioning in midbrain sections (**Fig. 3B**). To validate GRAB-DA and GRAB-ACh expression and fiber placement in DLS and DMS, we used an anti-GFP antibody (1:1000, Invitrogen #A11122) and anti-RFP antibody (1:1000) and Alexa 488 and 594 conjugated secondary antibodies (**Fig. 3C**; 1:500, Jackson Immunoresearch #711-546-152). We then mounted the sections with DAPI-containing mounting media (Southern Biotech #0100-20) and imaged fluorescence using a wide-field, slide-scanning fluorescence microscope (Keyence BZ-X800 or Leica Thunder).

## Results

### DMS DA and ACh dynamics under normal conditions

To investigate the temporal dynamics of striatal DA and ACh under normal conditions, we virally expressed fluorescent GRAB-DA and GRAB-ACh sensors in the dorsomedial striatum (DMS) of C57BL/6J mice. We then implanted an optical fiber into DMS injection site to enable dual-color fiber photometry in freely moving animals (**Fig. 1A, B**). As mice explored an open field arena, we recorded the fluorescence of each GRAB sensor during a 20-minute baseline period following vehicle injection, and for an additional 40 minutes following treatment with either the dopamine releasing drug amphetamine or the acetylcholinesterase inhibitor donepezil.

First, we examined the relationship between DA and ACh during the vehicle-treatment period. On average, DA and ACh fluctuations were anti-correlated with DA events followed by phasic decreases in ACh and ACh events followed by phasic decreases in DA (**Fig. 1C, D)**. Across the entire vehicle-treatment period, ACh and DA maintained an anti-correlated relationship with a slight temporal offset, consistent with previous reports^14^ (**Fig. 1E**). On average, the peak amplitudes of ACh transients and pauses that followed these transients were larger when a DA transient co-occurred within 2 s after the ACh transient peak (**Fig. 1F–K**), suggesting possible DAergic modulation. Finally, we found that ACh dynamics spon- taneously oscillated at of frequency of 1 Hz, consistent with previous reports^14^.

### DMS DA and ACh dynamics under hyperdopaminergic conditions

Next, we asked how these dynamics become altered under hyperdopaminergic conditions following amphetamine treatment (**Fig. 2A**). Systemic amphetamine treatment (2.5 mg·kg^-1^) increased bulk DA but minimally affected bulk ACh while slightly diminishing its anticorrelation to DA in the DMS (**Fig. 2B– E**). To validate this finding, we treated the mice with the acetylcholinesterase inhibitor donepezil and observed a bulk increase in GRAB-ACh fluorescence with an increase in bulk GRAB-DA fluorescence in the DMS (**Fig. 2F–H**).

**Figure 2.**
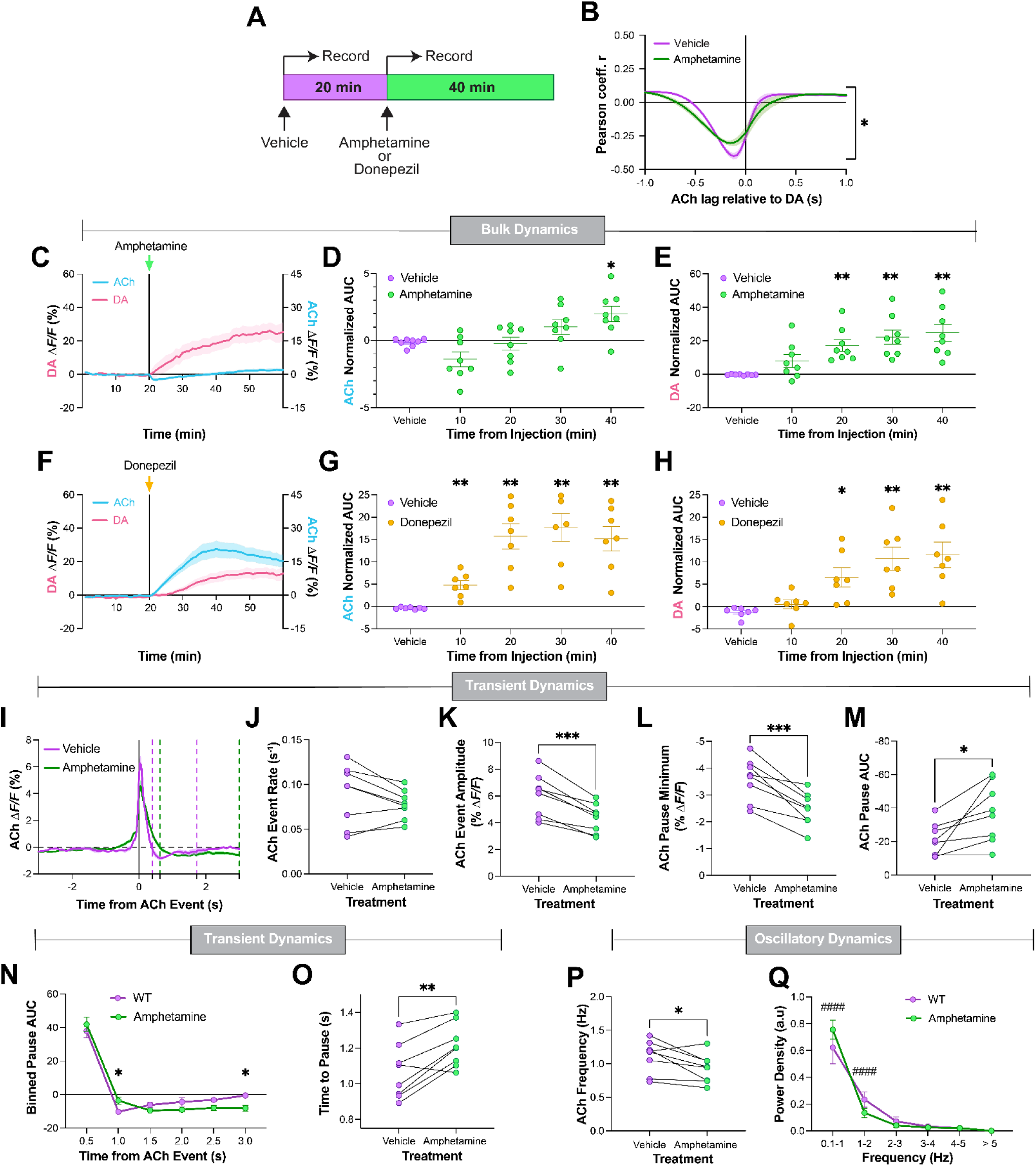
Amphetamine treatment effects on striatal DA and ACh dynamics. **(A)** Time course of photometry recordings and drug treatments in the open field. We administered vehicle at t = 0 min, rec- orded fluorescence for 20 min, injected amphetamine (2.5 mg·kg^-1^) or donepezil (2 mg·kg^-1^), and recorded fluorescence for 40 min. **(B)** Mean ± s.e.m. cross-correlation between ACh and DA signals following vehicle and amphetamine treatment. (\**P* < 0.05; two-way ANOVA effect of treatment) **(C)** Mean ± s.e.m. bulk GRAB-DA and GRAB-ACh fluorescence following vehicle and amphetamine treatment, normalized to the vehicle treatment period. **(D, E)** Mean ± s.e.m. AUC for bulk changes in GRAB-ACh, **D**, GRAB- DA, **E**, in 10-min bins following vehicle and amphetamine treatment, computed from vehicle-normalized Δ*F/F* traces (\**P* < 0.05 and \*\**P* < 0.01 compared to vehicle treatment; One-way ANOVA with Dunnett multiple comparisons). **(F–H)** Data corresponding to **C–E** following donepezil, rather than amphetamine treatment. **(I)** Mean ± s.e.m. GRAB-ACh event-triggered fluorescence following vehicle and ampheta- mine treatment. **(J–M)** Mean GRAB-ACh event rates, **J**, and event peak amplitudes, **K**, post-event pause minima, **L**, and post-event pause AUC, **M**, following vehicle and amphetamine treatment (\**P* < 0.05 and \*\*\**P* < 0.001; paired t-test). **(N)** Mean ± s.e.m. AUC of the GRAB-ACh pause after events in 0.5-second bins (\**P* < 0.05; two-way ANOVA with Holm Sidak multiple comparisons between treatment groups at indicated bins). **(O)** Mean time from event peak to the minimum value of the pause in GRAB-ACh fluo- rescence trace (\*\**P* < 0.01; paired t-test). **(P)** Mean ± s.e.m. weighted mean of GRAB-ACh fluorescence FFT frequency of GRAB-ACh following vehicle and amphetamine treatment (\**P* < 0.05; paired t-test). **(R)** Mean ± s.e.m. power spectral density of the GRAB-ACh in 1-Hz bins (two-way ANOVA with Holm Sidak multiple comparisons; ^####^*P* < 10^-4^). Data in **B–Q** are from *N* = 8 control mice.

**Figure 3.**
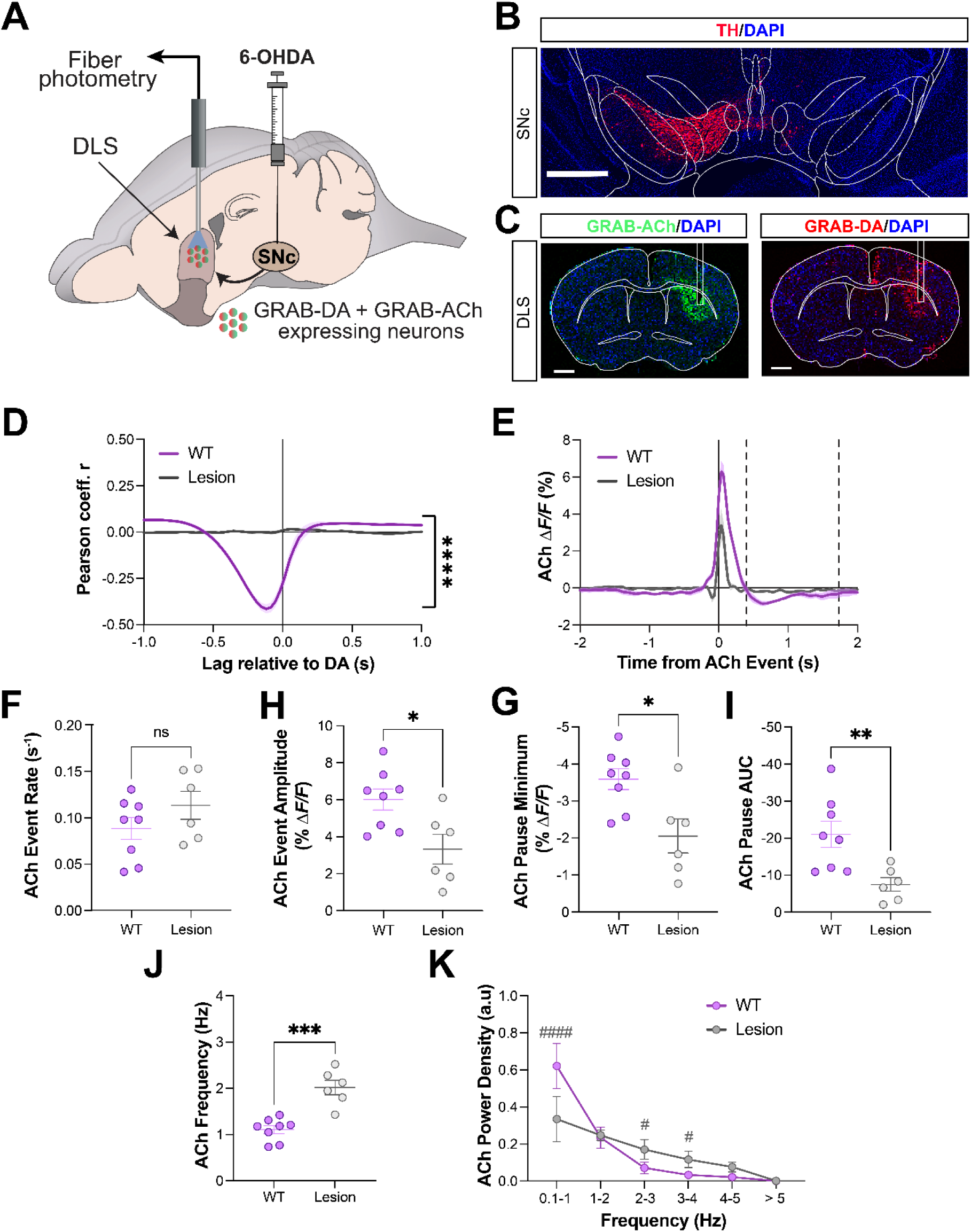
Dopamine depletion transforms striatal ACh dynamics. **(A)** We used an fiber optic to meas- ure GRAB-DA and GRAB-ACh fluorescence in the dopamine-depleted DLS of hemiparkinsonian mice injected with 6-OHDA in the SNc. **(B, C)** Tyrosine hydroxylase (TH) expression in the midbrain and GRAB-ACh and GRAB-DA expression in the DLS of a representative 6-OHDA-lesioned mouse (scale bar: 1 mm). **(D)** Mean ± s.e.m. cross-correlation of GRAB-ACh to GRAB-DA fluorescence in the DMS of un-lesioned (WT) and DLS of lesioned mice (\*\*\*\**P* < 10^-4^; two-way ANOVA main effect of group). **(E)** Mean ± s.e.m. GRAB-ACh event-triggered fluorescence aligned in WT and lesioned mice. **(F–I)** Mean GRAB-ACh event rates, **F**, and event peak amplitudes, **G**, post-event pause minima, **H**, and post- event pause AUC, **I**, following vehicle and amphetamine treatment (\**P* < 0.05 and \*\**P* < 0.01; paired t- test). **(J)** Mean ± s.e.m. weighted mean of GRAB-ACh fluorescence FFT frequency of GRAB-ACh in WT and lesioned mice (\*\*\**P* < 0.001; unpaired t-test) **(K)** Mean ± s.e.m. power spectral density of GRAB-ACh fluorescence in 1-Hz bins (^#^*P* < 0.05 and ^####^*P* < 10^-4^; two-way ANOVA with Holm Sidak multiple com- parisons). Data in **D–L** are from *N* = 8 WT and *N* = 6 lesioned mice.

For our remaining analyses, we focused on amphetamine’s effects on ACh dynamics in the DMS of these control mice. Amphetamine treatment did not affect the rate but did reduce the peak amplitude of ACh event transients (**Fig. 2I–K**). Amphetamine treatment also shallowed but prolonged the post-event pause in ACh signaling (**Fig. 2L–O**) and reduced the frequency of spontaneous ACh fluctuations^14^, altering their power density distribution (**Fig. 2P, Q**). These results reveal how a pharmacologically hyperdopaminergic state alters the bulk, transient, and oscillatory dynamics of ACh in the DMS of control mice.

### Effects of unilateral 6-OHDA lesion on DLS ACh dynamics

Next, we asked how ACh dynamics become altered under conditions modeling PD. To do this, we expressed GRAB-DA and GRAB-ACh sensors in the dorsolateral striatum (DLS) of C57BL/6J mice, unilaterally infused the neurotoxin 6-OHDA into the SNc projecting to the virus-injected hemisphere, and implanted a fiber optic probe to record DA and ACh dynamics during open field exploration (**Fig. 3A**). At the end of our recording experiments, we histologically verified SNc DA neurons lesions, virus expression, and fiber optic placement in the DLS (**Fig. 3B, C**).

Following 6-OHDA lesion, there was a complete loss of the normal anti-correlation between DA and ACh in the lesioned hemisphere, consistent with the loss of DA signaling (**Fig. 3D**). Compared to in the DMS of control mice, the rate of ACh transients in the DLS of lesioned mice was unchanged, but the peak amplitude and post-event pause of ACh transients were significantly reduced (**Fig. 3E–I**). Spectral analysis of spontaneous GRAB-ACh fluorescence fluctuations revealed a shift in the predominant signaling frequency band of from ∼1 Hz in the DMS of control mice to ∼2 Hz in the DLS of lesioned mice, an effect driven by decreased power density below 1 Hz and increased power density from 2–4 Hz (**Fig. 3J, K**). These findings reveal the multi-faceted transformation in DLS ACh dynamics following the loss of SNc dopamine neurons.

### Effects of L-DOPA on parkinsonian ACh dynamics

To evaluate how L-DOPA treatment affects these dynamics and whether this changes over the course of LID induction, we recorded GRAB-DA and GRAB-ACh in 6-OHDA lesioned mice following repeated L-DOPA injections. Specifically, two weeks after the 6-OHDA lesion, every other day we administered two initial doses of L-DOPA (6 mg·kg^−1^), then three doses of L-DOPA (12 mg·kg^−1^) to induce LID, followed by a final post LID induction L-DOPA dose (6 mg·kg^−1^) (**Fig. 4A**). This escalating dosing schedule has been shown to reliably induce LID in mice^21^. In each session we recorded GRAB-ACh and GRAB-DA dynamics for 20-min following vehicle treatment then for 40-min following L-DOPA treatment (**Fig. 4B**). During this LID induction period, we recorded following injections #1, 3, 4, and 6 (**Fig. 4A**). After LID induction, we performed an extended recording following a final L-DOPA dose (12 mg·kg^−1^) encompassing the transition to the L-DOPA OFF state—we recorded for 20 min following vehicle treatment and then for 180-min recording following L-DOPA treatment (t = 40–220 min after L-DOPA) (**Fig. 4A, C**).

**Figure 4.**
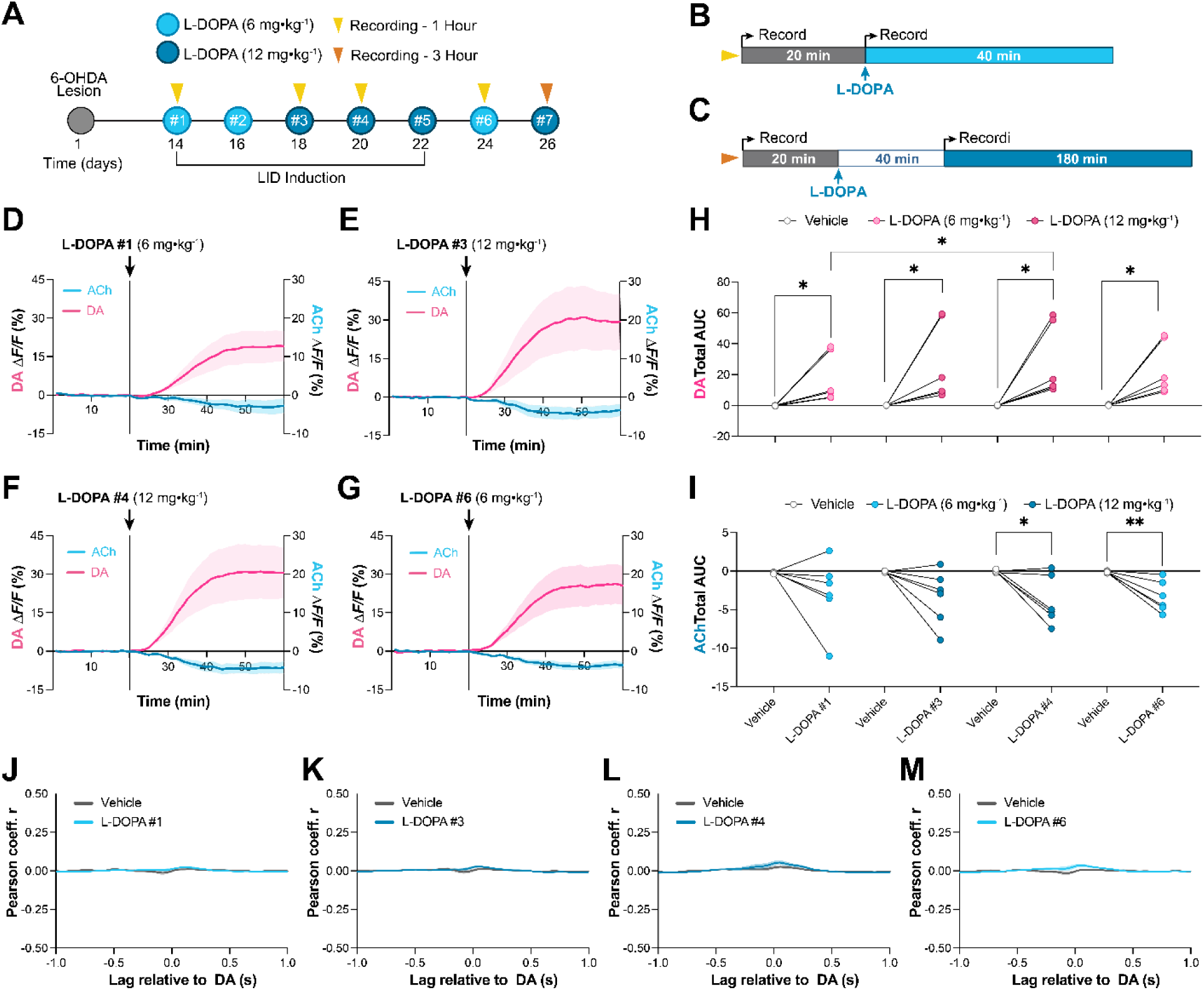
L-DOPA treatment and LID induction alter bulk DA and ACh dynamics. **(A)** Experimental protocol to measure the effects of L-DOPA on DA and ACh dynamics over the course of LID induction. **(B)** Time course for the first 4 photometry recordings and drug treatments in the open field. We administered vehicle at t = 0 min, recorded fluorescence for 20 min, injected L-DOPA (6 or 12 mg·kg^-1^), and recorded fluorescence for 40 min. **(C)** Time course for the final photometry recording and drug treatment in the open field. We administered vehicle at t = 0 min, recorded fluorescence for 20 min, injected L-DOPA (12 mg·kg^-1^), waited 40 minutes, and recorded fluorescence (toggling the recording on/off every 5 min) for 180 min. **(D–G)** Mean ± s.e.m. bulk GRAB-DA and GRAB-ACh fluorescence following vehicle and the corresponding L-DOPA treatments (6 or 12 mg·kg^-1^) in our experimental protocol, **A**, normalized to the vehicle treatment period. **(H, I)** Mean ± s.e.m. AUC for bulk changes in GRAB-DA, **H**, and GRAB-ACh, **I**, fluorescence following vehicle and L-DOPA treatment (\**P* < 0.05 and \*\**P* < 0.01; paired t-test). **(J–M)** Mean ± s.e.m. cross-correlation between GRAB-DA and GRAB-ACh fluorescence following vehicle and L-DOPA treatments (P > 0.05; two-way ANOVA main effect of treatment). Data in **D–M** are from *N* = 6 lesioned mice.

*Bulk DA and ACh dynamics and cross-correlation:* Treatment with both doses of L-DOPA significantly increased bulk DA in the DLS following each injection, and injection #4 (12 mg·kg^−1^) evoked greater DA signaling than injection #1 (6 mg·kg^−1^) (**Fig. 4D–H**). Although the effects of the initial L-DOPA injections were not significant, injection #4 (12 mg·kg^−1^) during LID induction and injection #6 (6 mg·kg^−1^) after LID induction significantly reduced bulk ACh signaling in the DLS (**Fig. 4F**, **G**, & **I**). Despite modulating bulk DA and ACh in the DLS, L-DOPA treatment failed to restore the pronounced anticorrelated relationship between DA and ACh observed under normal conditions in the DMS (**Fig. 2B**), as indicated by Pearson correlation coefficients near zero across all temporal lags (**Fig. 4J–M**).

*ACh transient dynamics*: L-DOPA treatment during (12 mg·kg^−1^) and after (6 mg·kg^−1^) LID induction reduced both the event rate of ACh transients (**Fig. 5A–E**). Compared to the DMS of WT mice, L-DOPA treatment further reduced the peak amplitudes of ACh events in the DLS of 6-OHDA lesioned mice (**Fig. 5D, E**). L-DOPA treatment also diminished the depth of post-transient pauses in ACh signaling, which were already shallower in the DLS of 6-OHDA lesioned mice compared to the DMS of WT mice (**Fig. 5F, G**). Compared to the ACh pause minima, the overall reduced AUC of post-transient ACh pauses was unaffected by L-DOPA except for the final 6 mg·kg^−1^ injection after LID induction, which further reduced the AUC of ACh pauses (**Fig. 5H, I**).

**Figure 5.**
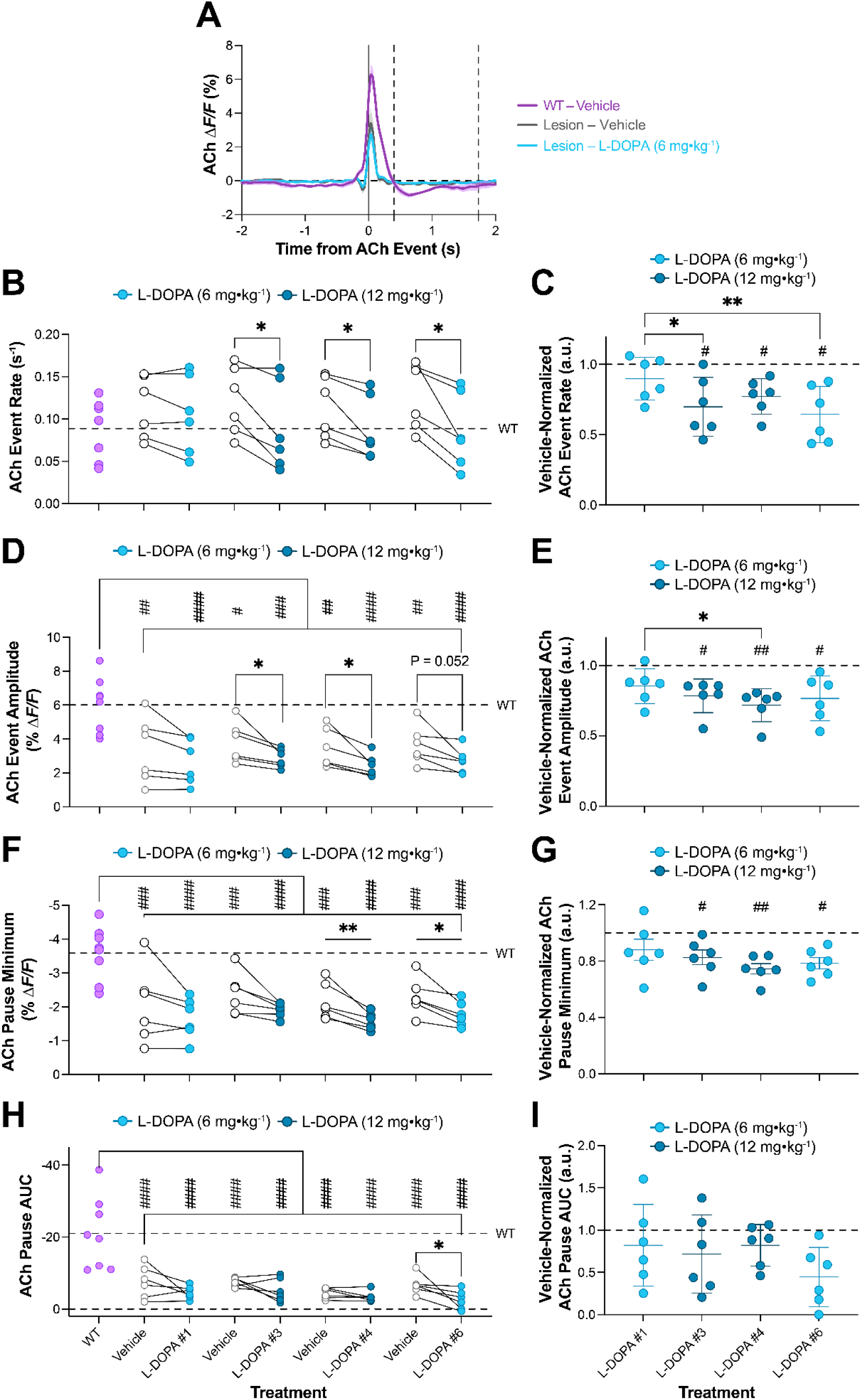
L-DOPA does not normalize ACh transient dynamics. **(A)** Mean ± s.e.m. GRAB-ACh event- triggered fluorescence in WT and lesioned mice following vehicle and L-DOPA treatment #1 (6 mg·kg^-1^). **(B)** Mean rates of GRAB-ACh events in WT mice and lesioned mice following vehicle and L-DOPA treatments (\**P* < 0.05; paired t-test). (**C**) Same as in **B** but normalized to values following vehicle treatment (\**P* < 0.05 and \*\**P* < 0.01 comparing normalized L-DOPA doses; ^#^*P* < 0.05 compared to normalized vehicle values; one-way ANOVA with Dunnett multiple comparisons). (**D**) Mean event-triggered GRAB- ACh fluorescence peak amplitudes in WT and lesioned mice following vehicle and L-DOPA treatments (\**P* < 0.05 and \*\**P* < 0.01; paired t-test; ^#^*P* < 0.05, ^##^*P* < 0.01, ^###^*P* < 0.001, and ^####^*P* < 10^-4^; one-way ANOVA with Dunnett multiple comparisons to WT values). (**E**) Same as in **D** but normalized to values following vehicle treatment (\**P* < 0.05 comparing normalized L-DOPA doses; ^#^*P* < 0.05 and ^##^*P* < 0.01 compared to normalized vehicle values; one-way ANOVA with Dunnett multiple comparisons). (**F**) Mean post-event GRAB-ACh pause minima in WT mice and lesioned mice following vehicle and L-DOPA treatments (\**P* < 0.05 and \*\**P* < 0.01; paired t-test; ^###^*P* < 0.001 and ^####^*P* < 10^-4^; one-way ANOVA with Dunnett multiple comparisons to WT values). (**G**) Same as in **F** but normalized to values following vehicle treatment (^#^*P* < 0.05 and ^##^*P* < 0.01 compared to normalized vehicle values; one-way ANOVA with Dun- nett multiple comparisons). (**H**) Mean post-event GRAB-ACh pause AUC in WT mice and lesioned mice following vehicle and L-DOPA treatments (\**P* < 0.05; paired t-test; ^###^*P* < 0.001 and ^####^*P* < 10^-4^; one- way ANOVA with Dunnett multiple comparisons to WT values). (**I**) Same as in **H** but normalized to values following vehicle treatment. All data are from *N* = 8 WT and *N* = 6 lesioned mice.

*ACh oscillatory dynamics*: L-DOPA treatment during (12 mg·kg^−1^) and after (6 mg·kg^−1^) LID induction reduced the prominent frequency of spontaneous ACh oscillations, though not to the level observed in the DMS of WT mice (**Fig. 6A, B**). Specifically, L-DOPA treatment increased the power density of ACh oscillations below 1 Hz and reduced the power density of ACh oscillations from 2–4 Hz, though the power densities in these ranges remained significantly different than the densities observed in the DMS of WT mice (**Fig. 6C, D**).

**Figure 6.**
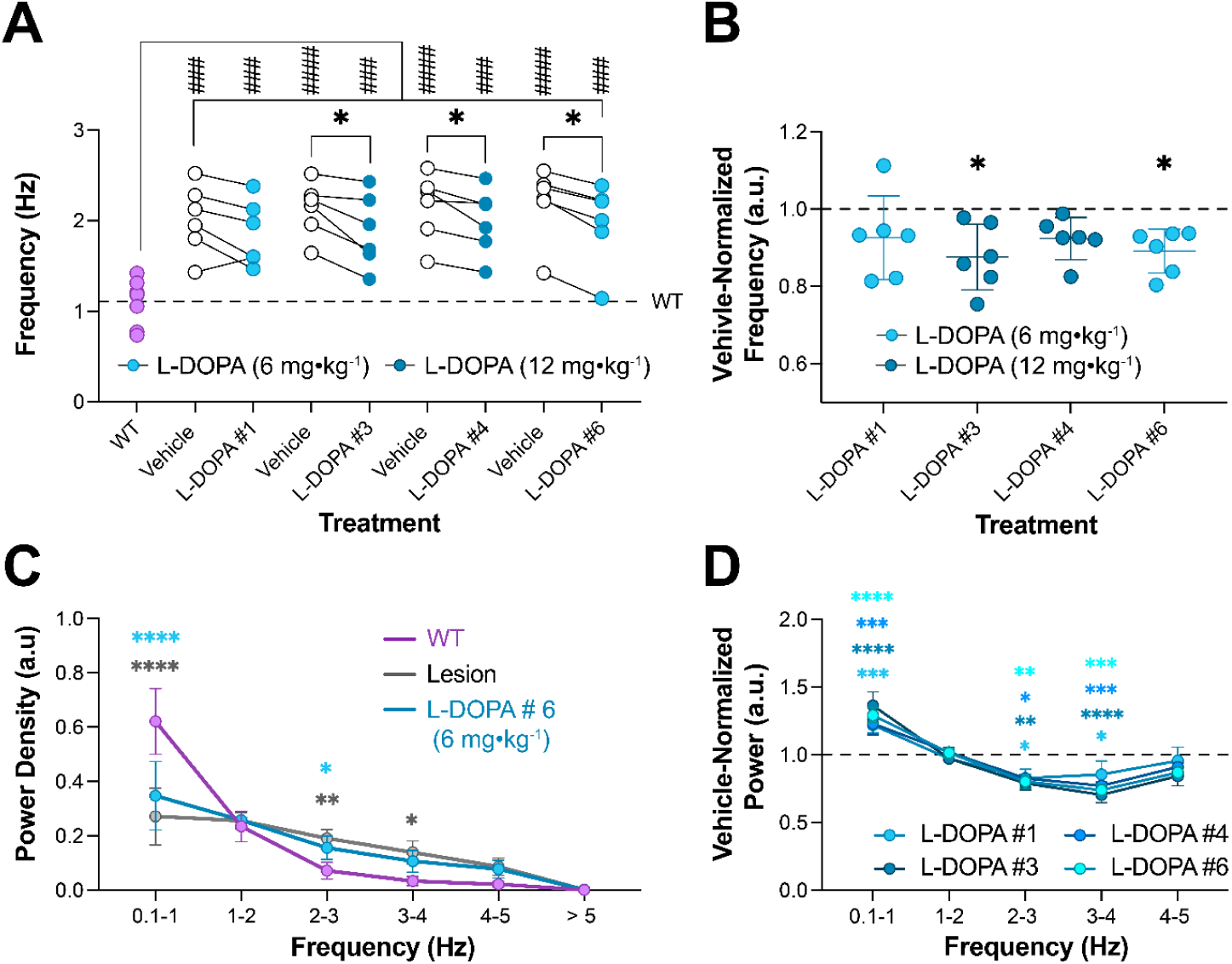
L-DOPA fails to normalize oscillatory ACh dynamics. **(A)** Weighted mean of fluorescence FFT frequency of GRAB-ACh in lesioned mice following vehicle and L-DOPA treatments (\**P* < 0.05; paired t-test compared to vehicle; ^###^*P* < 0.001 and ^####^*P* < 10^-4^; one-way ANOVA with Holm-Sidak multiple comparisons to WT values). **(B)** Weighted mean of fluorescence FFT frequency of GRAB-ACh signals in lesioned mice following L-DOPA treatments, normalized to values following vehicle treatments (\**P* < 0.05; one-way ANOVA with Dunnett multiple comparisons to normalized vehicle treatment values). **(C)** Mean ± s.e.m. power spectral density of GRAB-ACh fluorescence in 1-Hz bins for WT mice and lesioned mice following vehicle and L-DOPA treatment #6 (6 mg·kg^-1^) after LID induction (\**P* < 0.05, \*\**P* < 0.01, and \*\*\*\**P* < 10^-4^; two-way ANOVA with Holm-Sidak multiple comparisons to WT values). **(D)** Same as in **C,** for values after L-DOPA treatments, normalized to power values following vehicle treatments in lesioned mice (\**P* < 0.05, \*\**P* < 0.01 and \*\*\**P* < 0.001; two-way ANOVA with Holm-Sidak multiple comparisons to normalized vehicle values). All data are from *N* = 8 WT and *N* = 6 lesioned mice.

Altogether, these results demonstrate that despite increasing DA and reducing ACh signaling in the DLS, L-DOPA failed to normalize the temporal dynamics of ACh signaling. In particular, L-DOPA treatment failed to restore the normal anticorrelation between DA and ACh, it did not fully normalize the rate of ACh events and further exacerbated the reduced peak amplitude and diminished post-event pause of ACh events, and it did not fully reverse the heightened frequency of spontaneous ACh fluctuations following 6-OHDA lesion.

### ACh dynamics during transition to L-DOPA OFF state

To investigate the time-dependent alterations in cholinergic signaling during the transition to the L-DOPA OFF state, we analyzed the bulk, transient and oscillatory ACh dynamic in 1-hour bins, for 3 hours starting 40 minutes after a final L-DOPA (12 mg·kg^−1^; **Fig. 4C**). During the initial 40 minutes after L-DOPA treatment, GRAB-DA fluorescence was increased and GRAB-ACh fluorescence decreased (**Fig. S1A**). While bulk DA signaling progressively returned to baseline levels over the course of the three-hour recording, bulk ACh levels remained lower throughout the OFF-state transition (**Fig. S1B, C**). L-DOPA treatment initially reduced the rate of ACh events, an effect that waned over the three-hour recording (**Fig. S1D**). By contrast, L-DOPA treatment reduced the peak amplitude of ACh transients and the depth (but not overall AUC) of post-event pauses in ACh signaling, effects that lasted throughout the OFF-state transition (**Fig. S1E–G**). Finally, L-DOPA treatment initially reduced the frequency of spontaneous ACh oscillations, though this effect also went away later in the three-hour recording (**Fig. S1H**).

In summary, during the transition to the L-DOPA OFF-state, bulk DA signaling subsided, but bulk ACh signaling remained low. Although the rate of ACh events and frequency of ACh oscillations returned to pre-treatment levels during the OFF-state transition, the peak amplitude of ACh events and the depth of post-event pauses in ACh signaling remained lower throughout the transition.

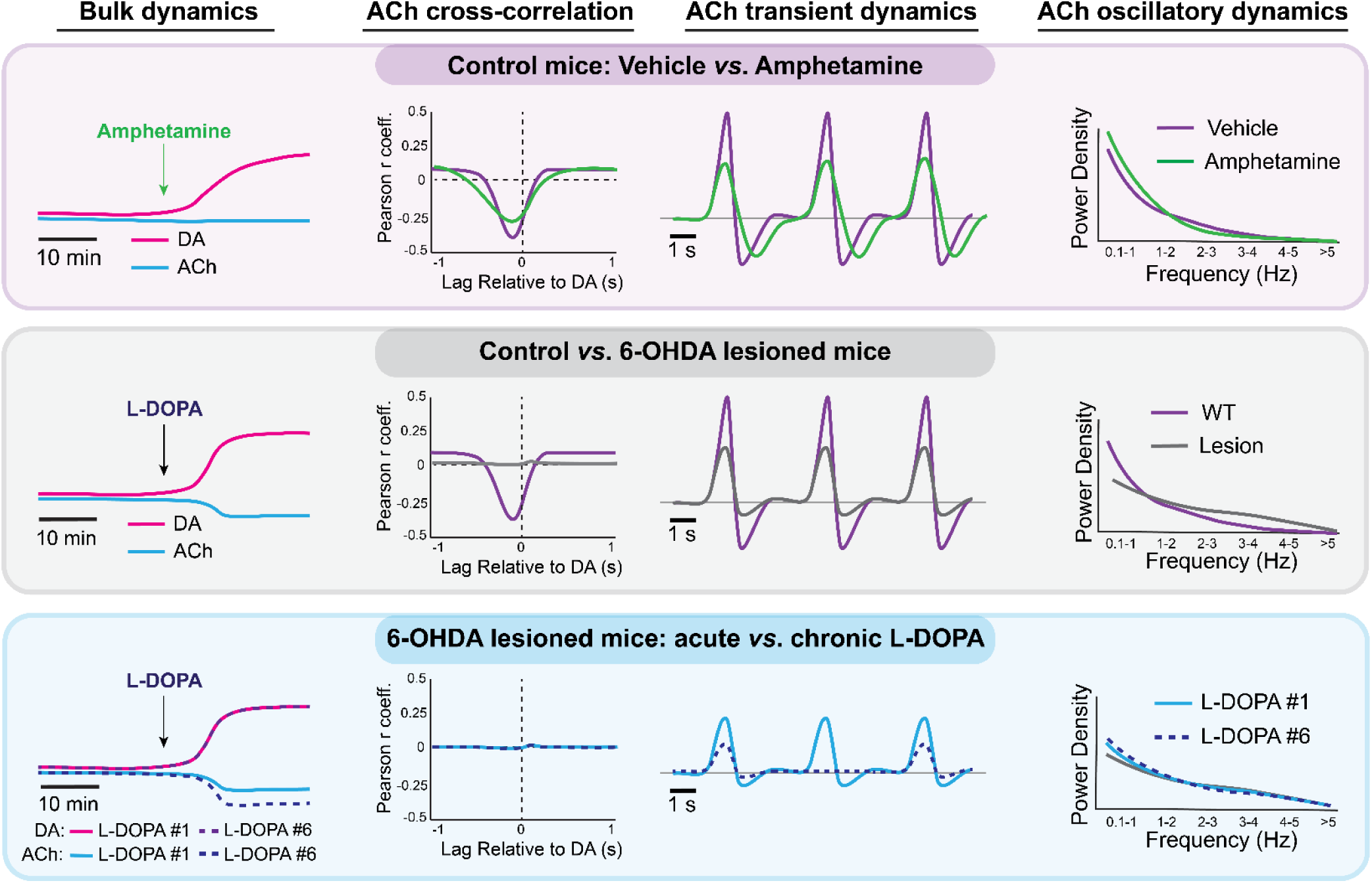

## Discussion

Here we used fiber photometry GRAB sensors for DA and ACh to determine how these neuromodulators become altered by dopamine under normal and parkinsonian conditions. Our work builds upon several recent studies to further elucidate how DA/ACh dynamics become altered in the context of PD and L-DOPA treatment^8,14,16^. As with our study, each of these studies hinge upon the advent of neuromodulator biosensors and the ability to simultaneously record sub-second DA and ACh fluctuations. Using these powerful tools, these studies collectively demonstrate that DAergic manipulations transform the bulk, transient, and oscillatory dynamics of ACh. Notably, most of the altered dynamics after DA cell loss were not normalized by L-DOPA treatment. These results advance our understanding of the normal interrelationship between DA and ACh and how this relationship may go awry in PD.

### DA and ACh dynamics in the DMS under normal conditions

Consistent with prior reports and classical models, ACh dynamics were temporally anti-correlated to DA in the DMS of C57BL/6J mice^14^ (**Fig. 1C–E**). Not only was fluorescence between GRAB-ACh and GRAB-DA anti-correlated, ACh event transients preceded a reduction in DA and vice versa. Consistent with this earlier study, we also found that ACh spontaneously fluctuates with prominent frequency range of 0.1–2 Hz (**Fig. 1L, M**). Although this ACh fluctuation has been shown to be driven by excitatory glutamatergic innervation (rather than DAergic innervation), local interaction in the striatum may still modulate these anti-correlated DA and ACh dynamics^6,13^. Consistent with this idea, a closer examination revealed that the co-occurrence of DA and ACh transients modulated the depth of the pause in ACh transmission following ACh transients (**Fig. 1F–K**). It has been argued that the activation of inhibitory D2 receptors in ACh interneurons may directly modulate ACh pauses^6^. The fact that the peak of ACh transients that preceded DA events were larger than ACh transients that did not may reflect greater synchronized ACh interneuron activity for these events, which could be responsible for facilitating DA release^22,23^.

### ACh dynamics in the DMS under hyperdopaminergic conditions in normal mice

Treating mice with amphetamine to induce a hyperdopaminergic state elevated DA signaling but less significantly modulated bulk ACh levels (**Fig. 2C–E**). Under these conditions, DA and ACh largely maintained their anti-correlated relationship, despite a modest, but significant reduction (**Fig. 2B**). Amphetamine treatment also reduced the peak amplitude of ACh events, prolonged the pause in ACh following these events, and reduced the frequency of spontaneous ACh fluctuations (**Fig. 2I–Q**).

Amphetamine increases extracellular DA levels by entering and blocking the dopamine transporter (DAT) in DA neuron terminals and displacing vesicular DA, causing it to undergo reverse transport through DAT^24,25^. Amphetamine’s multiplexed mechanism-of-action means that it can induce DA release independently of SNc DA neuron activity, but it may also diminish activity-dependent, pulsatile DA release due to vesicular DA occlusion^26^. By increasing basal DA and reducing synaptic vesicular DA, amphetamine may have reduced the signal-to-noise ratio of DA release dynamics, which may account for the modestly diminished anti-correlation between ACh and DA fluorescence.

An increase in bulk DA may also underlie the reduced peak amplitudes of ACh events and the prolongation of ACh pauses after ACh events. Consistent with this, signaling at D2Rs, which are expressed in ACh interneurons, has been shown to modulate the duration (and AUC) of ACh pauses in response to reward-predictive stimuli^6^. Although the effects of amphetamine treatment on the frequency of ACh fluctuations were modest, D2R activation may also have contributed to this change. For example, although not quantified in their study, DA receptor antagonists appear to cause an upward shift in the frequency power distribution of ACh fluctuations^14^. Therefore, in our study, D2R activation may have dampened ACh interneuron responsiveness to the excitatory afferents thought to drive these dynamics, though more experiments are necessary to support this conclusion.

### ACh dynamics in the DA-depleted DLS

Following 6-OHDA lesion of SNc DA neurons, GRAB-ACh fluorescence predictably had no anti-correlation to GRAB-DA fluorescence (**Fig. 3D**). Compared to data in the DMS of WT mice, ACh transients in the DA-depleted DLS had diminished amplitudes, and the post-transient pauses in ACh were abolished (**Fig. 3E–I**). ACh dynamics also fluctuated at a higher frequency in the DA-depleted DLS (**Fig. 3J, K**).

One limitation of our study is that our control data were from the DMS, while the DA-depleted data were from the DLS. This makes it difficult to conclude that any differences between the WT and Lesion experimental groups were not simply innate differences between the DMS and DLS. However, although there is evidence for sub-regional differences in striatal ACh responses to reward delivery^14^, the described differences are more qualitative than quantitative (*i.e*., the presence peaks and pauses in ACh, rather than their precise magnitudes). Moreover, another study reported quantitatively similar dynamics between spontaneous ACh fluctuations in the DLS and DMS^6^. While further systematic studies are warranted, we found the DMS data from control mice to be a useful benchmark for understanding how DA depletion transforms ACh dynamics in DLS, notwithstanding this apparent caveat.

The diminution of post-transient pauses in ACh as well as the increased frequency of spontaneous ACh fluctuations in the DA-depleted DLS are directionally opposite to the effects of increasing dopamine via amphetamine treatment in the DMS. Here again, it is also possible that the loss of D2R signaling ACh interneurons altered these metrics in the DA-depleted DLS. It was previously reported spontaneous fluctuations in ACh persist after DA depletion, but increase in amplitude, and frequency^14^ (consistent with our finding). Taken together with the amphetamine treatment data, these findings suggest that DA may modulate these oscillatory dynamics, even if it is not their primary driver.

It is less clear why both amphetamine treatment (**Fig. 2K**) and DA depletion (**Fig. 3H**) reduced the amplitude of ACh events. One possible explanation is that DA depletion in our study chronically elevated ACh levels in the DLS, thereby diminishing the dynamic range of pulsatile ACh event transients. By contrast, we did not observe changes in bulk ACh levels in the DMS following acute amphetamine treatment, suggesting the reduction in the amplitude we observed following amphetamine treatment likely resulted from the increase, rather than decrease in DA, likely through D2Rs. Although we cannot measure basal ACh levels using GRAB-ACh, this interpretation is consistent with previous reports of increased ACh tone of the striatum in PD animal models^3,27^.

### L-DOPA treatment effects on ACh dynamics in the DA-depleted DLS

L-DOPA predictably increased DA and, after repeated treatments, diminished ACh levels in the DA-depleted DLS (**Fig. 4D–I**). Despite re-balancing DA and ACh, L-DOPA treatment did not re-establish the anti-correlated relationship between GRAB-ACh and GRAB-DA fluorescence (**Fig. 4J–M**). L-DOPA reduced the rate of ACh event transients after repeated treatments and diminished the peak amplitude of these events, which were already lower than in the DMS of WT mice (**Fig. 5B–E**). L-DOPA did not restore the depth or AUC of the pauses in ACh signaling after event transients; in fact, repeated treatments further diminished these pause metrics (**Fig. 5F–I**). While L-DOPA did partially reduce the frequency of spontaneous ACh fluctuations, it did not do so to the levels in the DMS of WT mice (**Fig. 6**). Finally, during the transition to a L-DOPA OFF state, DA returned to baseline levels while ACh levels remained reduced, and the peak amplitude of ACh event transients and the magnitude of pauses remained diminished (**Fig. S1**).

One study recently linked reductions in bulk ACh following L-DOPA treatment in DA-depleted striatum to the activation of D2R in ACh interneurons and to D1R signaling in GABAergic interneuron that modulate ACh interneurons^8^. Therefore, as with amphetamine treatment, it is likely that L-DOPA treatment leads to DA release that engages, in this case hypersensitized DA receptors^28^, in the DA-depleted DLS to directly or indirectly suppress ACh transmission. The emergent effects of L-DOPA on ACh after repeated treatments and the emergence of LID may reflect the further sensitization of the DA receptor signaling^29^. For example, we found that L-DOPA also had emergent effects on the peak amplitude of ACh events and the oscillatory dynamics of ACh, which may also reflect changes in DA receptor sensitivity following 6-OHDA lesions and repeated L-DOPA. The facts that L-DOPA treatment failed to normalize the anti-correlation between DA and ACh and the post-event pause in ACh transmission likely reflects an uncoupling between DA and ACh transmission due to non-physiological DA release from non-DA neurons in the 6-OHDA lesioned hemisphere.

### Implications for PD

Under normal conditions, the functional role of fast, anti-correlated dynamics between DA and ACh is not fully understood, but their coordination is thought to be critical for gating synaptic plasticity in D1- and D2-SPNs^4^. For example, DA facilitates LTP in D1-SPNs, while inhibitory M4 receptor activation can oppose this process^4,5,7^. Thus, a coordinated rise in DA and decline in ACh, as observed under normal conditions, may be permissive for DA-dependent changes in synaptic strength in SPNs. Not only did DA depletion perturb these anti-correlated dynamics, but it also diminished the peak amplitude of ACh transients and the minima of subsequent pauses in ACh transmission. These dynamics may also be important for modulating activity or synaptic strength in SPNs. Despite increasing DA and reducing ACh at large, the fact that L-DOPA treatment failed to normalize or even exacerbated the effects of DA depletion on the transient and oscillatory dynamics of ACh poses a complication for PD. This loss of dynamic ACh signaling to coordinate DA-dependent plasticity in the striatum may uncouple DA transmission from behavioral relevance, resulting in aberrant changes in synaptic strength and neural activity that further disrupt movement.

Although it is unclear how to restore the ACh dynamics that are lost by DA depletion, add-on treatments like M4 ACh receptor positive allosteric modulators (PAMs) have shown promise for LID in animal models^5^. M4 receptors are also autoreceptors in ACh interneurons, so augmenting their activity could feasibly augment ACh post-transient pauses or modulate the frequency of spontaneous ACh fluctuations. Further studies are necessary to determine how these or other treatments targeted to ACh interneurons will help determine whether these dynamics can be targeted and whether doing so has therapeutic potential.

---

**Figure S1.**
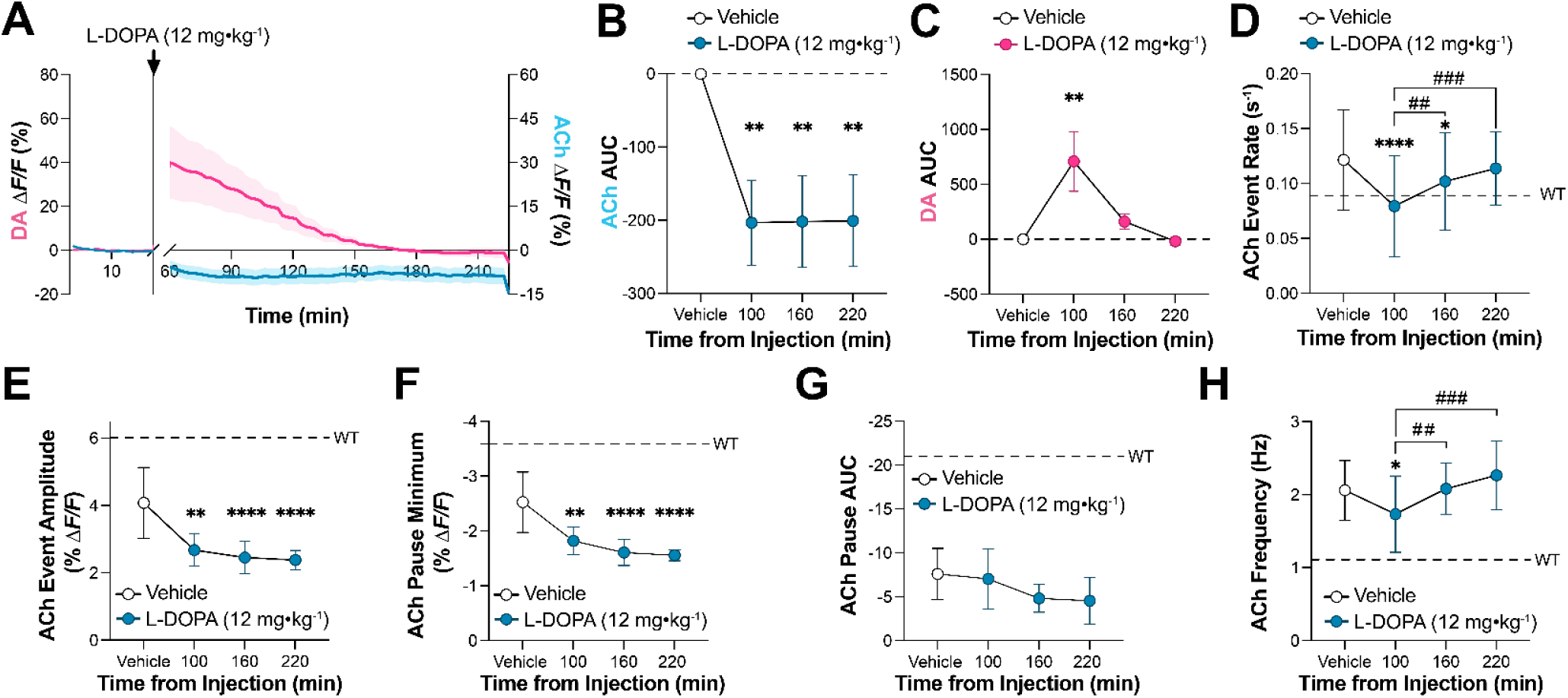
DA and ACh dynamics during transition to L-DOPA OFF State. **(A)** Mean ± s.e.m. bulk GRAB-DA and GRAB-ACh fluorescence following the final treatment (#7) of lesioned mice with vehicle and L-DOPA (12 mg·kg^-1^) using the extended imaging protocol in Fig. 4C. Data are normalized to the last 40% of the vehicle treatment period. **(B, C)** Mean ± s.e.m. AUC for bulk changes in GRAB-ACh, **B**, and GRAB-DA, **C**, fluorescence following vehicle and L-DOPA treatment in 60-min bins (\*\**P* < 0.01; one-way ANOVA with Holm Sidak multiple comparisons vehicle treatment). **(D–G)** Mean ± s.e.m. GRAB-ACh event rate, **D**, and event peak amplitude, **E**, post-event pause minimum, **F**, and post-event pause AUC, **G**, in lesioned mice following vehicle and L-DOPA treatment (\*\**P* < 0.01 and \*\*\*\**P* < 10^-4^ compared to vehicle treatment; ^##^*P* < 0.01 and ^###^*P* < 0.001 comparing post L-DOPA treatment periods; one-way ANOVA with Holm Sidak multiple comparisons). **(H)** Mean ± s.e.m. weighted mean of GRAB-ACh fluorescence FFT frequency following vehicle and L-DOPA treatment (\**P* < 0.05 and compared to vehicle treatment; ^##^*P* < 0.01 and ^###^*P* < 0.001 comparing post L-DOPA treatment periods; one-way ANOVA with Holm Sidak multiple comparisons). All data from *N* = 6 lesioned mice.

## References

1. Hoover, D.B., Muth, E.A., and Jacobowitz, D.M. (1978). A mapping of the distribution of acetycholine, choline acetyltransferase and acetylcholinesterase in discrete areas of rat brain. Brain Res. 153, 295–306. 10.1016/0006-8993(78)90408-0.

2. Versteeg, D.H., Van Der Gugten, J., De Jong, W., and Palkovits, M. (1976). Regional concentrations of noradrenaline and dopamine in rat brain. Brain Res. 113, 563–574. 10.1016/0006-8993(76)90057-3.

3. Pisani, A., Bernardi, G., Ding, J., and Surmeier, D.J. (2007). Re-emergence of striatal cholinergic interneurons in movement disorders. Trends Neurosci. 30, 545–553. 10.1016/j.tins.2007.07.008.

4. Wang, Z., Kai, L., Day, M., Ronesi, J., Yin, H.H., Ding, J., Tkatch, T., Lovinger, D.M., and Surmeier, D.J. (2006). Dopaminergic control of corticostriatal long-term synaptic depression in medium spiny neurons is mediated by cholinergic interneurons. Neuron 50, 443–452. 10.1016/j.neuron.2006.04.010.

5. Shen, W., Plotkin, J.L., Francardo, V., Ko, W.K., Xie, Z., Li, Q., Fieblinger, T., Wess, J., Neubig, R.R., Lindsley, C.W., et al. (2015). M4 Muscarinic Receptor Signaling Ameliorates Striatal Plasticity Deficits in Models of L-DOPA-Induced Dyskinesia. Neuron 88, 762–773. 10.1016/j.neuron.2015.10.039.

6. Martyniuk, K.M., Torres-Herraez, A., Lowes, D.C., Rubinstein, M., Labouesse, M.A., and Kellendonk, C. (2022). Dopamine D2Rs coordinate cue-evoked changes in striatal acetylcholine levels. eLife 11. 10.7554/eLife.76111.

7. Shen, W., Flajolet, M., Greengard, P., and Surmeier, D.J. (2008). Dichotomous dopaminergic control of striatal synaptic plasticity. Science 321, 848–851. 10.1126/science.1160575.

8. Li, H., Chen, Z., Tan, Y., Luo, H., Lu, C., Gao, C., Shen, X., Cai, F., Hu, J., and Chen, S. (2024). Enhancing striatal acetylcholine facilitates dopamine release and striatal output in parkinsonian mice. Cell Biosci 14, 146. 10.1186/s13578-024-01328-z.

9. Zhang, Y., Rózsa, M., Liang, Y., Bushey, D., Wei, Z., Zheng, J., Reep, D., Broussard, G.J., Tsang, A., Tsegaye, G., et al. (2023). Fast and sensitive GCaMP calcium indicators for imaging neural populations. Nature 615, 884–891. 10.1038/s41586-023-05828-9.

10. Patriarchi, T., Cho, J.R., Merten, K., Howe, M.W., Marley, A., Xiong, W.H., Folk, R.W., Broussard, G.J., Liang, R., Jang, M.J., et al. (2018). Ultrafast neuronal imaging of dopamine dynamics with designed genetically encoded sensors. Science 360. 10.1126/science.aat4422.

11. Howe, M., Ridouh, I., Allegra Mascaro, A.L., Larios, A., Azcorra, M., and Dombeck, D.A. (2019). Coordination of rapid cholinergic and dopaminergic signaling in striatum during spontaneous movement. Elife 8. 10.7554/eLife.44903.

12. Chantranupong, L., Beron, C.C., Zimmer, J.A., Wen, M.J., Wang, W., and Sabatini, B.L. (2023). Dopamine and glutamate regulate striatal acetylcholine in decision-making. Nature 621, 577–585. 10.1038/s41586-023-06492-9.

13. Gallo, E.F., Greenwald, J., Yeisley, J., Teboul, E., Martyniuk, K.M., Villarin, J.M., Li, Y., Javitch, J.A., Balsam, P.D., and Kellendonk, C. (2022). Dopamine D2 receptors modulate the cholinergic pause and inhibitory learning. Mol. Psychiatry 27, 1502–1514. 10.1038/s41380-021-01364-y.

14. Krok, A.C., Maltese, M., Mistry, P., Miao, X., Li, Y., and Tritsch, N.X. (2023). Intrinsic dopamine and acetylcholine dynamics in the striatum of mice. Nature 621, 543–549. 10.1038/s41586-023-05995-9.

15. Matityahu, L., Gilin, N., Sarpong, G.A., Atamna, Y., Tiroshi, L., Tritsch, N.X., Wickens, J.R., and Goldberg, J.A. (2023). Acetylcholine waves and dopamine release in the striatum. Nature communications 14, 6852. 10.1038/s41467-023-42311-5.

16. Atamna, Y., Tiroshi, L., Wattad, N., Gilin, N., Gilad, S., Berkowitz, N., and Goldberg, J.A. (2025). Levodopa Disrupts Activity Patterns and Encoding of Movement in Striatal Cholinergic Interneurons of Behaving Mice. Mov Disord. 10.1002/mds.70018.

17. Hamid, A.A., Frank, M.J., and Moore, C.I. (2021). Wave-like dopamine dynamics as a mechanism for spatiotemporal credit assignment. Cell 184, 2733–2749.e2716. 10.1016/j.cell.2021.03.046.

18. Barbeau, A. (1962). The pathogenesis of Parkinson’s disease: a new hypothesis. Can Med Assoc J 87, 802–807.

19. Ztaou, S., Maurice, N., Camon, J., Guiraudie-Capraz, G., Kerkerian-Le Goff, L., Beurrier, C., Liberge, M., and Amalric, M. (2016). Involvement of Striatal Cholinergic Interneurons and M1 and M4 Muscarinic Receptors in Motor Symptoms of Parkinson’s Disease. J Neurosci 36, 9161–9172. 10.1523/jneurosci.0873-16.2016.

20. Corder, G., Ahanonu, B., Grewe, B.F., Wang, D., Schnitzer, M.J., and Scherrer, G. (2019). An amygdalar neural ensemble that encodes the unpleasantness of pain. Science 363, 276–281. 10.1126/science.aap8586.

21. Fieblinger, T., Graves, S.M., Sebel, L.E., Alcacer, C., Plotkin, J.L., Gertler, T.S., Chan, C.S., Heiman, M., Greengard, P., Cenci, M.A., and Surmeier, D.J. (2014). Cell type-specific plasticity of striatal projection neurons in parkinsonism and L-DOPA-induced dyskinesia. Nature communications 5, 5316. 10.1038/ncomms6316.

22. Zhang, Y.F., Reynolds, J.N.J., and Cragg, S.J. (2018). Pauses in Cholinergic Interneuron Activity Are Driven by Excitatory Input and Delayed Rectification, with Dopamine Modulation. Neuron 98, 918–925.e913. 10.1016/j.neuron.2018.04.027.

23. Brimblecombe, K.R., Threlfell, S., Dautan, D., Kosillo, P., Mena-Segovia, J., and Cragg, S.J. (2018). Targeted Activation of Cholinergic Interneurons Accounts for the Modulation of Dopamine by Striatal Nicotinic Receptors. eNeuro 5. 10.1523/eneuro.0397-17.2018.

24. Sulzer, D., Chen, T.K., Lau, Y.Y., Kristensen, H., Rayport, S., and Ewing, A. (1995). Amphetamine redistributes dopamine from synaptic vesicles to the cytosol and promotes reverse transport. J Neurosci 15, 4102–4108. 10.1523/jneurosci.15-05-04102.1995.

25. Fleckenstein, A.E., Volz, T.J., Riddle, E.L., Gibb, J.W., and Hanson, G.R. (2007). New insights into the mechanism of action of amphetamines. Annu Rev Pharmacol Toxicol 47, 681–698. 10.1146/annurev.pharmtox.47.120505.105140.

26. Schmitz, Y., Lee, C.J., Schmauss, C., Gonon, F., and Sulzer, D. (2001). Amphetamine distorts stimulation-dependent dopamine overflow: effects on D2 autoreceptors, transporters, and synaptic vesicle stores. J Neurosci 21, 5916–5924. 10.1523/jneurosci.21-16-05916.2001.

27. Ding, J., Guzman, J.N., Tkatch, T., Chen, S., Goldberg, J.A., Ebert, P.J., Levitt, P., Wilson, C.J., Hamm, H.E., and Surmeier, D.J. (2006). RGS4-dependent attenuation of M4 autoreceptor function in striatal cholinergic interneurons following dopamine depletion. Nat. Neurosci. 9, 832–842. 10.1038/nn1700.

28. Gerfen, C.R. (2000). Molecular effects of dopamine on striatal-projection pathways. Trends Neurosci. 23, S64–70. 10.1016/s1471-1931(00)00019-7.

29. Feyder, M., Bonito-Oliva, A., and Fisone, G. (2011). L-DOPA-Induced Dyskinesia and Abnormal Signaling in Striatal Medium Spiny Neurons: Focus on Dopamine D1 Receptor-Mediated Transmission. Front Behav Neurosci 5, 71. 10.3389/fnbeh.2011.00071.

